# Cross-modal temporal biases emerge during early sensitive periods

**DOI:** 10.1101/839514

**Authors:** Stephanie Badde, Pia Ley, Siddhart S Rajendran, Idris Shareef, Ramesh Kekunnaya, Brigitte Röder

## Abstract

Human perception features stable biases, such as perceiving visual events as later than synchronous auditory events. The origin of such perceptual biases is unknown, they could be innate or shaped by sensory experience during a sensitive period. To investigate the role of sensory experience, we tested whether a congenital, transient loss of vision, caused by bilateral dense cataracts, has sustained effects on the ability to order events spatio-temporally within and across sensory modalities. Most strikingly, individuals with reversed congenital cataracts showed a bias towards perceiving visual stimuli as occurring earlier than auditory (Exp. 1) and tactile (Exp. 2) stimuli. In contrast, both normally sighted controls and individuals who could see at birth but developed cataracts during childhood reported the typical bias of perceiving vision as delayed compared to audition. Thus, we provide strong evidence that cross-modal temporal perceptual biases depend on sensory experience and emerge during an early sensitive period.

## Introduction

In every moment, a multitude of information reaches our brain through the different senses. These sensory inputs need to be separated, ordered in space and time to derive a coherent representation of the environment. Yet, the perception of temporal order is seldom veridical. Reports illustrating the subjectivity of cross-modal temporal perception date back to 18^th^ and 19^th^ century astronomers; small but stable individual biases in the perceived timing of visual and auditory events caused significant differences in the measurements of stellar transit times and subsequent scientific disputes. These early reports inspired the pioneering work of Wilhelm Wundt and promoted the role of perceptual biases as a major but still unresolved topic in experimental psychology ^1,2^.

Determining the spatio-temporal order of events across sensory modalities poses an especially difficult challenge, as information arriving through different senses travels at different speeds – in the environment and within the nervous system ^3^. However, the typical bias towards perceiving vision as delayed compared to audition cannot be explained by these physical and physiological speed differences ^4–6^.

The perceived temporal order of visual-auditory events can be transiently shifted by exposing humans to a series of asynchronous stimulus pairs with a constant lag between vision and audition ^7,8^. Yet, such recalibration effects quickly vanish and in the long-term cross-modal temporal biases are highly stable within individuals ^1,6^. This co-existence of short-term plasticity and long-term stability re-emphasizes the question that already troubled scientists 250 years ago: why does the brain not learn to compensate for such perceptual biases?

Given their long-term stability, biases in cross-modal temporal perception could either be innate, inherent to the structure of the underlying neural mechanisms, or emerge based on sensory experience potentially during a sensitive period of development. The ability to optimally order spatially and temporally distinct events across sensory modalities develops only in late childhood and overall later than within one sensory modality ^9^. This late maturation points toward the possibility that cross-modal spatio-temporal biases are shaped by sensory experience accumulated during childhood. The role of early sensory experience for the genesis of perceptual biases is extremely difficult to address in humans; only naturally altered developmental sensory environments open a window into the influence of experience on perceptual development. We tested the hypothesis of a sensitive period for the emergence of cross-modal perceptual biases by measuring the ability to order events spatio-temporally across vision, audition, and touch in individuals born with dense, bilateral cataracts whose sight was restored 6 to 168 months after birth.

## Results

Thirteen individuals with a history of transient, congenital bilateral, dense cataracts (CC) participated in two spatial temporal order judgement tasks, ten in a visual-auditory task (Exp. 1) and ten in a visual-tactile task (Exp. 2). To test for the role of vision during infancy as well as to control for the role of persisting visual impairments, sixteen individuals, nine per experiment, who underwent surgery for cataracts which had developed after birth during childhood (DC) served as controls, additionally to age-matched typically sighted individuals (MCC and MDC). In every trial, two successive stimuli were presented, one in each hemifield. Visual-auditory and visual-tactile stimulus pairs were randomly interleaved with unimodal stimulus pairs. Participants reported the side of the first stimulus irrespective of its modality ^5^. We predicted preferential processing of and consequently an increased bias toward the auditory and tactile modality as well as a lower visual and cross-modal spatio-temporal resolution in the CC-group.

Most strikingly and contrary to our predictions, CC-individuals showed a bias toward perceiving visual stimuli as occurring earlier than auditory (Fig. 1A; bias CC-group, Expt. 1, χ^2^(1) = 4.31, *p* = 0.038; see Supplementary File 1A for full statistical models) and tactile stimuli (Fig. 1C; bias CC-group, Expt. 2, χ^2^(1) = 12.46, *p* < 0.001), respectively. This bias did not significantly correlate with the duration of their visual deprivation, the time passed since sight was restored, and their restored visual acuity (see Figure 2 and Supplementary File 1B). CC-individuals’ bias towards perceiving visual stimuli as earlier stood in marked contrast to the bias observed for their matched controls (bias group-difference CC-vs. MCC-group, Expt. 1, visual-auditory, χ^2^(1) = 8.48, *p* = 0.004; Expt. 2, visual-tactile, χ^2^(1) = 5.31, *p* = 0.021) who perceived auditory stimuli as occurring earlier than visual stimuli (bias MCC-group, Expt. 1, visual-auditory, χ^2^(1) = 4.18, *p* = 0.041) and did not show a significant bias for visual-tactile comparisons (bias MCC-group, Expt. 2, visual-tactile, χ^2^(1) = 0.15, *p* = 0.701). DC-individuals showed a typical bias towards perceiving visual stimuli as occurring later than auditory and tactile stimuli (bias DC-group, Expt. 1, visual-auditory, χ^2^(1) = 7.14, *p* = 0.008; Expt. 2, visual-tactile, χ^2^(1) = 8.86, *p* = 0.003). In fact, their spatio-temporal bias toward perceiving tactile stimuli as earlier than visual stimuli even exceeded the bias of their matched controls (bias group-difference DC-vs. MDC-group, Expt. 1, visual-auditory, χ^2^(1) = 1.84, *p* = 0.175; Expt. 2, visual-tactile, χ^2^(1) = 8.08, *p* = 0.004; bias MDC-group, Expt. 1, visual-auditory, χ^2^(1) = 0.83, *p* = 0.362; Expt. 2, visual-tactile, χ^2^(1) = 0.85, *p* = 0.356).

**Figure 1.**
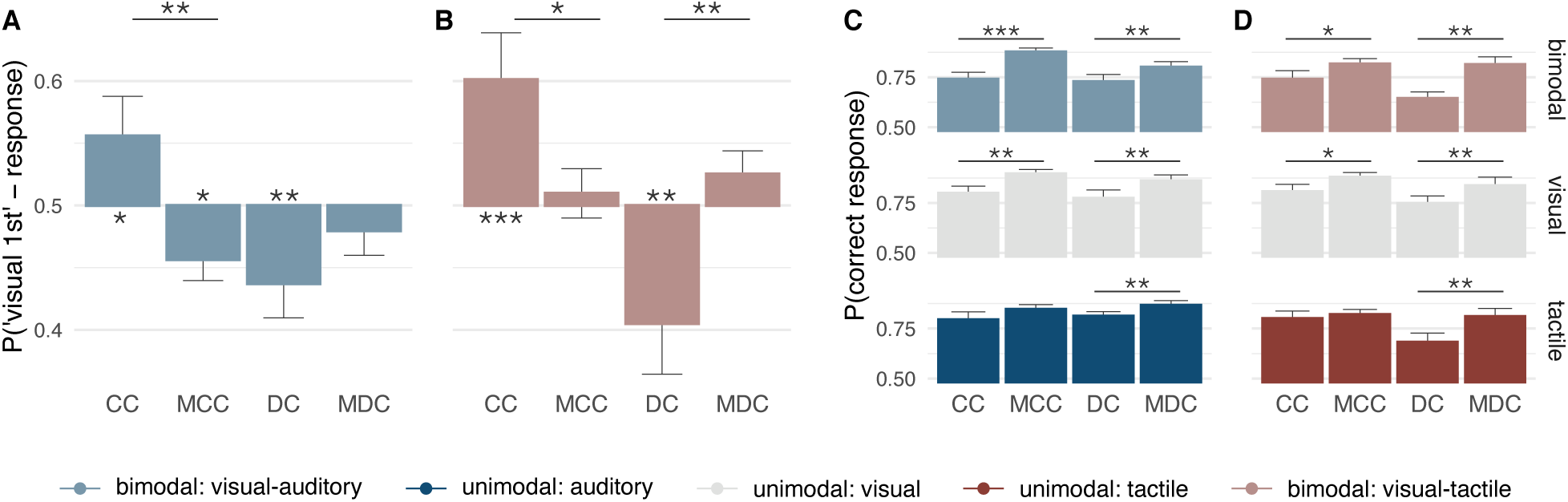
Effects of transient visual deprivation on spatio-temporal order biases and resolution within and across vision, audition, and touch. Thirteen individuals with a history of congenital, bilateral dense cataracts (CC) and sixteen individuals whose reversed cataracts had developed later in life (DC) as well as age-, gender-, and handedness-matched typically-developed individuals (MCC, MDC) took part in the study (see Table 1 for details about the samples). Participants judged the spatio-temporal order of two successive stimuli – one presented in each hemifield – by indicating the location of the first stimulus. In Expt. 1, visual (grey), auditory (dark blue), and visual-auditory (light blue) stimuli were presented, in Expt. 2, visual, tactile (dark red), and visual-tactile (light red) stimuli. (A, B) Spatio-temporal biases: group average probabilities of perceiving the visual stimulus as preceding the auditory (A) or the tactile (B) stimulus in bimodal trials. Probabilities are shown relative to chance level. (C, D) Spatio-temporal resolution: group average probabilities of correct responses separately for each modality condition. Error bars indicate standard errors of the mean.

**Figure 2.**
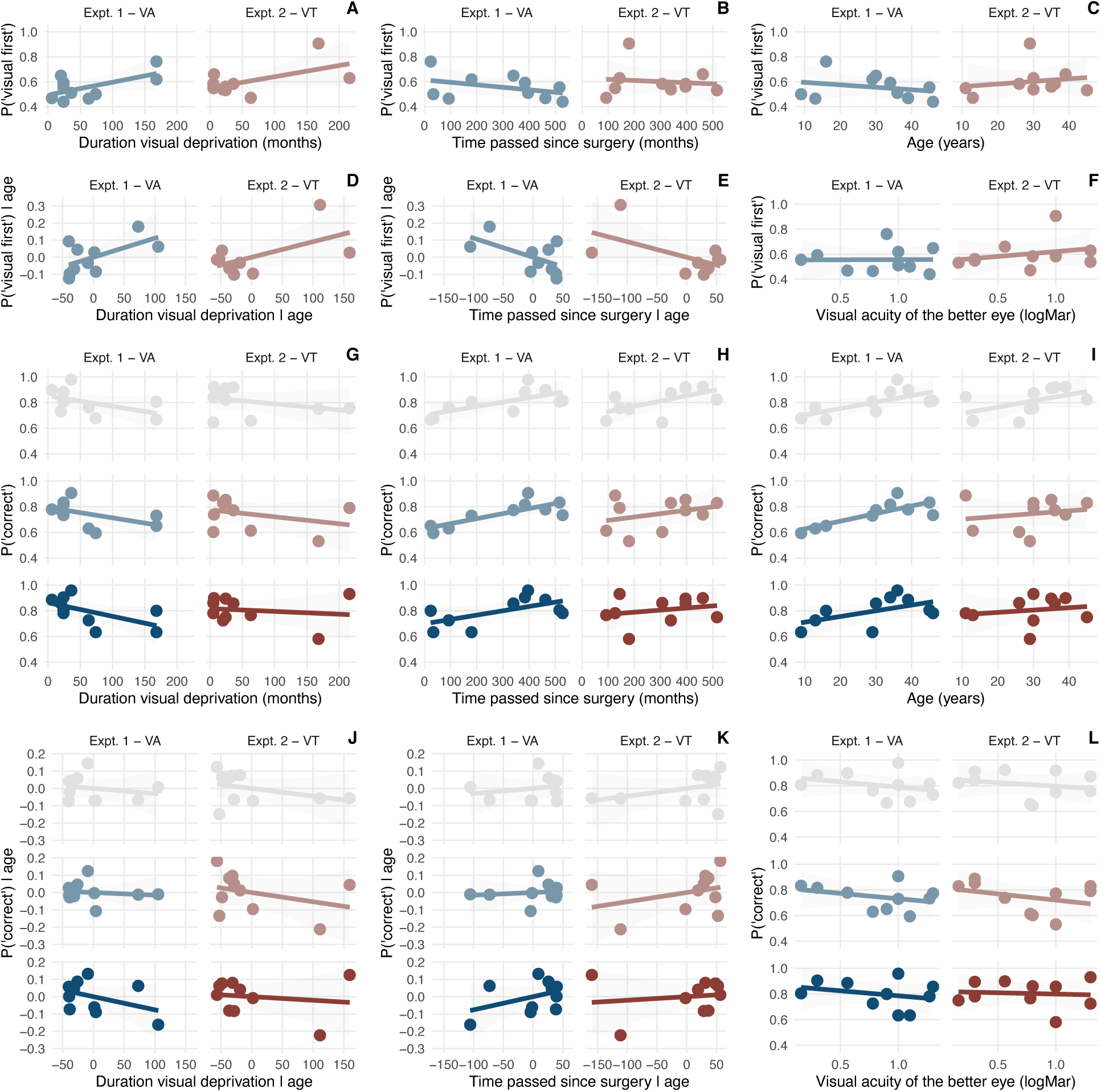
Correlations between measures of spatio-temporal perception and key variables of CC-individuals’ visual history. (A-C,F) Correlation between CC-individuals’ cross-modal bias measured as the proportion of visual first responses and (A) the duration of their visual deprivation, (B) the time passed between surgery and testing, (C) their age, and (E) the visual acuity of the better eye. (D,E) Partial correlations between the proportion of visual first responses and (D) the duration of their visual deprivation as well as (E) the time passed between surgery and testing corrected for age, to this aim, each variable refers to the residuals of a linear model predicting the respective variable from age. (G-L) Correlations and partial correlations between CC-individuals’ spatio-temporal resolution measured as the proportion of correct responses and the same variables as before separately for each modality condition and experiment.

CC-individuals’ visual and cross-modal spatio-temporal resolution (Fig. 1B,D) was reduced compared to that of their controls (resolution group-difference CC-group vs. MCC-group, Expt. 1, visual, χ^2^(1) = 9.68, *p* = 0.002; Expt. 1, visual-auditory, χ^2^(1) = 17.88, *p* < 0.001; Expt. 2, visual, χ^2^(1) = 4.72, *p* = 0.030; Expt. 2, visual-tactile, χ^2^(1) = 4.03, *p* = 0.045), but no significant difference between the CC-group and their controls emerged for the auditory and tactile modalities, (auditory resolution group-difference CC-group vs. MCC-group, χ^2^(1) = 1.42, *p* = 0.233; tactile resolution group difference, χ^2^(1) = 0.11, *p* = 0.742). DC-individuals’ spatio-temporal resolution (Fig. 1B,D) was lower than the resolution of their controls independent of modality condition (resolution group-difference DC-group vs. MDC-group, Expt. 1, χ^2^(1) = 4.98, *p* = 0.026; Expt. 2, χ^2^(1) = 11.99, *p* = 0.001). CC-individuals’ spatio-temporal resolution did not significantly correlate with their key medical history (see Supplementary File 1B; Figure 2).

**Table 1.** Sample characteristics

|  | N | Sex | Handedness | Visual acuity of the better eye | Age at testing | Age at surgery | Time period between surgery and testing |
| --- | --- | --- | --- | --- | --- | --- | --- |
| CC – Expt. 1 visual-auditory | 10 | 8 males | 10 right-handed | 0.16 – 1.3 logMar, mean 0.84 logMar | 9 – 46 years, mean 30 years | 6 – 168 months, mean 60 months | 24 – 528 months, mean 296 months |
| DC – Expt. 1 visual-auditory | 9 | 7 males | 9 right-handed | -0.5 – 0.8 logMar, mean 0.25 logMar | 8 – 19 years, mean 13 years | 74 – 183 months, mean 120 months | 12 – 66 months, mean 30 months |
| CC – Expt. 2 visual-tactile | 10 | 10 males | 10 right-handed | 0.16 – 1.3 logMar, mean 0.75 logMar | 11 – 45 years, mean 30 years | 5 – 216 months, mean 57 months | 93 – 516 months, mean 296 months |
| DC – Expt. 2 visual-tactile | 9 | 6 males | 9 right-handed | 0 – 0.5 logMar, mean 0.22 logMar | 9 – 19 years, mean 14 years | 30 – 183 months, mean 102 months | 13 – 174 months, mean 63 months |

## Discussion

We investigated whether cross-modal perceptual biases are innate or acquired during a sensitive phase in early childhood by testing sight-recovery individuals with a history of congenital visual loss. In two spatio-temporal order judgement tasks (visual-auditory and visual-tactile) CC-individuals reported visual stimuli as earlier than both auditory and tactile stimuli, exhibiting a reversed cross-modal bias compared to their controls and sight-recovery individuals whose cataracts had developed later. Moreover, the ability to determine the spatio-temporal order of separate events across different sensory modalities and within vision was reduced after a transient phase of visual loss at any time during childhood. These results, for the first time, demonstrate that cross-modal perceptual biases are not innate but rather acquired during a sensitive period.

At first glance, CC-individuals’ bias to perceive visual events as earlier than either auditory or tactile events seems to indicate that visual stimuli are processed faster than auditory and tactile stimuli following transient, congenital visual deprivation. However, CC-individuals’ spatio-temporal resolution revealed a temporal processing impairment for vision but not for audition and touch. Consistently, event-related potentials have provided no evidence for earlier visually-evoked brain activity in CC-individuals ^10,11^ and behavioral studies have demonstrated no visual advantage in reaction times to simple visual stimuli ^12,13^. Moreover, reduced visual contrast – typical for cataract-reversal individuals – is associated with delayed responses of the visual system and lower visual temporal sensitivity ^14,15^. Thus, there is currently no evidence indicating accelerated processing of visual information following congenital visual deprivation. Indeed, accelerated visual processing in the CC-group would be counterintuitive, given that these individuals had no reliable visual input for extensive periods after birth. Alternatively, biases in temporal order perception can be induced by allocating attention unequally across modalities ^16–19^. However, attention has been ruled out as a cause of persistent biases in cross-modal spatio-temporal processing ^6^. Moreover, it is not obvious why CC- and DC-individuals would entertain opposing attentional foci given that both groups experienced transient severe visual impairment and suffer from remaining visual acuity impairments. In sum, the diverging cross-modal temporal biases observed here between groups unlikely reflect differences in the speed of processing or attentional effects but rather represent a difference in the genuine perceptual biases of humans born with and without vision.

CC-individuals’ surprising bias toward perceiving visual stimuli as occurring earlier than auditory and tactile stimuli is consistent with exposure to lagging visual stimuli while the cataracts were still present: Residual light perception, which exists even in the presence of dense cataracts, likely has been sluggish due to reduced retinal transduction rates ^20^ and suppression of visual cortex activity in the context of cross-modal stimulation ^21^. Thus, cataract-reversal individuals likely were exposed to a consistent delay of vision compared to audition and touch before cataract reversal surgery, resulting in a reversed visual-auditory and visual-tactile bias after cataracts were removed. Crucially, the CC-individuals were exposed to these altered sensory environments from birth on. Studies in owls have demonstrated that atypical multisensory experience due to prism-altered vision leads to permanent structural differences in the mapping of auditory and visual spatial representations in the juvenile brain ^22^ but not in the adult brain ^23^. Thus, we suggest that the reversed bias exhibited by CC-individuals results from structural differences elicited by atypical cross-modal temporal experience after birth. In sum, stable cross-modal temporal biases ^5,6^ might exist because the brain optimizes cross-modal temporal perception ^24^ during a sensitive period and as a consequence establishes a setpoint for future recalibration.

The finding, that CC-individuals but not DC-individuals showed a reversed visual-auditory temporal bias provides strong evidence that perceptual biases are shaped by sensory experience during early childhood. In contrast to CC-individuals, DC-individuals had encountered temporally aligned cross-modal stimuli after birth, which we suggest had enabled them to develop a typical bias. It has been shown in cats that even minimal visual experience prior to visual deprivation allows for a typical development of cortico-cortical connections ^25^. The sensitive period for cross-modal temporal perception might span the first six months of life, the minimal duration of deprived vision in our CC-group. A previous study tested CC-individuals whose vision was restored within early infancy (4 months of age on average) and their matched controls on simultaneity judgments of spatially aligned visual-auditory and visual-tactile stimulus pairs but did not report a reversed cross-modal bias ^26^. However, since simultaneity judgments of spatially aligned cross-modal stimuli are less sensitive to biases than the spatial temporal order judgements we used, it remains open whether these CC-individuals showed no reversed cross-modal bias because their sight was restored before the sensitive period was over. In sum, our results provide strong evidence for a sensitive period for the emergence of cross-modal spatio-temporal biases during human ontogeny.

Both cataract reversal groups exhibited a lower spatio-temporal resolution, indicative of an increased temporal uncertainty, than their controls in visual and cross-modal contexts. This general disadvantage suggests a strong dependence of cross-modal temporal ordering on the visual sense that might be related to the spatial nature of our task. Moreover, the finding that both cataract groups exhibited a lower temporal resolution but only the CC-group an altered cross-modal bias strongly suggests that cross-modal spatio-temporal biases and resolution are dissociable processes. Moreover, the conjunction of CC- and DC-individuals’ reduced resolution might point towards a long sensitive period for the development of spatio-temporal sensory resolution which would be compatible with the protracted developmental time course of multisensory temporal processing ^9,27^. Yet, persisting visual impairments might have contributed to the reduced spatio-temporal resolution of both cataract-reversal groups.

Furthermore, the present finding of increased temporal uncertainty could explain why recent studies have persistently found altered multisensory integration following congenital, transient periods of visual deprivation ^26,28,29^. A higher spatio-temporal uncertainty predicts wider temporal integration windows for simple, spatially-aligned stimuli ^26^, and at the same time hinders the detection of temporal correlations ^30^ between more complex signals such as speech stimuli ^28,29^.

In conclusion, congenital but not late transient visual deprivation was associated with a bias towards perceiving visual events as earlier than auditory or tactile events, suggesting an early sensitive period for the development of perceptual biases.

## Methods

### Participants

The sample of the visual-auditory experiment (Expt. 1) comprised 10 individuals who were born with bilateral dense cataracts (CC) and whose vision was restored later in life (for details see Table 1) and 9 individuals with transient, bilateral cataracts which had developed during childhood (DC). The sample tested in the visual-tactile experiment (Expt. 2) comprised 10 CC- and 9 DC-individuals. For each CC- and DC-participant an age-, gender- and handedness-matched control participant was recruited. Seven CC- and 2 DC-individuals as well as 13 control participants took part in both experiments. The majority of participants with a history of cataracts were recruited and tested at the LV Prasad Eye Institute in Hyderabad, India. Three CC-individuals and all control participants were recruited and tested at the University of Hamburg, Germany. The presence of congenital cataracts was affirmed by medical records. Since cataracts were sometimes diagnosed at a progressed age, additional criteria such as presence of nystagmus, the density of the lenticular opacity, the lack of fundus visibility prior to surgery, a family history of congenital cataracts, and parents’ reports were employed to confirm the onset of the cataract. Data of five additional participants were excluded from all analyses because the onset of the cataract remained unclear (2 participants, 1 took part in the visual-auditory experiment and 1 in the visual-tactile experiment), the time period between surgery and testing was shorter than 12 months (1 participant from the CC-group who took part in both experiments and 1 participant from the DC-group who took part in the visual-auditory experiment), or because additional neurological problems were suggested by the medical records (1 participant from the CC-group who took part in both experiments). Data of two further participants (1 CC- and 1 DC-individual) were excluded because they performed at chance level in the visual-auditory experiment. All excluded data are shown in the supplementary information (Fig. 1 – Figure Supplement 1) and are in accordance with the results presented in the main text. Adult participants and legal guardians of minors were reimbursed for travel expenses, accommodation, and absence from work, if applicable; adult participants tested in Hamburg received a small monetary compensation or course credit. Children received a small present. All participants or, if applicable, their legal guardian, provided written informed consent before beginning the experiment. The study was conducted in accordance with the ethical guidelines of the Declaration of Helsinki and was approved by the ethical board of the German Psychological Society as well as the local ethical committee of the Hyderabad Eye Research Foundation.

### Apparatus and Stimuli

Participants sat at a table, facing two speakers, positioned at 14° visual angle (15 cm at 60 cm distance) to the left and to the right of the participant’s midline. Three LEDs were mounted on top of each speaker. In the visual-tactile experiment, custom-made, noise-attenuated tactile stimulators were attached to the dorsal sides of both index fingers. For stimulation, the LEDs emitted red light, the speakers played white noise, and the tactile stimulators vibrated at a frequency of 100 Hz. Each stimulus lasted 15 ms, independent of modality. All three LEDs were used for cataract-reversal participants, but only one LED for typically sighted participants to roughly compensate for persistent visual impairments in cataract-reversal participants. To rule out that typically sighted participants perceived vision as delayed due to the lower number of LEDs, we tested 5 additional typically sighted participants (all female and right-handed, 23-50 years old, mean age 34 years) in the visual-auditory experiment while using all three LEDs. These participants too showed a significant bias towards perceiving vision as delayed (*t*(4)=6.36, *p*=0.003, Fig. 1 – Figure Supplement 2). Constant white noise was presented from a centrally located speaker to mask residual noise produced by the tactile stimulators. During the experiment, participants fixated a mark placed centrally between the loudspeakers and rested both hands on buttons aligned with the loudspeakers (visual-auditory experiment) or both feet on footpedals (visual-tactile experiment). Younger participants sometimes experienced problems activating the response devices in a controlled manner. These participants (visual-auditory experiment: 1 CC-individual; visual-tactile experiment: 2 DC-individuals) and their controls responded by waving one hand and the experimenter entered the response. The experiment was controlled by Presentation (Version 17.1.05, Neurobehavioral Systems, Inc., Berkeley, CA, www.neurobs.com), which recorded responses and interfaced with custom-built hardware to drive the stimulators.

### Task, Procedure, and Design

In each trial, two stimuli were presented in close succession; one stimulus in each hemifield. Participants indicated at which side they perceived the first stimulus. Responses had to be withheld until the second stimulus had been presented. Response times were not restricted, and the next trial started 2 s after the response had been registered.

The modality of the stimulus presented at either side (visual or auditory, Expt. 1; visual or tactile, Expt. 2) and the stimulus onset asynchrony (SOA; ±30, ±90, ±135, ±400 ms, with negative SOAs indicating ‘left side first’-stimulus pairs) of the two stimuli varied pseudo-randomly across trials. Each of the 32 stimulus conditions (2 modalities x 2 sides x 8 SOAs) was repeated 10 times; the 320 trials were divided into 10 blocks. Participants additionally completed ten practice trials with an SOA of ±400 ms at the beginning of the experiment. If necessary, the practice trials were repeated until participants felt confident about the task. In the visual-tactile experiment, a subsample of participants was additionally tested while holding the hands crossed (data not reported here). Participants were encouraged to take breaks in between blocks. Some of the cataract-reversal participants did not complete the full experiment, mostly due to time constraints. Except for practice trials, participants did not receive feedback.

### Data Analysis

Data and analysis scripts are made available online ^31^. Trials with reaction times shorter than 100 ms and more than 2.5 standard deviations above the participant’s mean reaction time (RT) were excluded from the analysis (2.1% of trials; responses entered by the experimenter were not filtered).

Each participants’ data were split according to the modality of the stimulus pair (visual-visual, auditory-auditory, or visual-auditory for Expt. 1; visual-visual, tactile-tactile, or visual-tactile for Expt. 2). Participants’ left-right responses in bimodal trials were transformed into binary ‘visual first’ – values.

To test for temporal order biases toward one modality, we conducted a hierarchical logistic regression on single-trial ‘visual first’-values with group as predictor. As planned comparisons, we first calculated pairwise contrasts comparing each cataract-reversal group with its matched control group and second estimated fixed contrasts separately for each group to evaluate whether the probability to perceive the visual stimulus before the auditory or tactile stimulus significantly differed from chance level.

To analyze the spatio-temporal resolution across groups and modality conditions, we conducted a hierarchical logistic regression on single trial accuracy values using group and modality as predictors. To resolve interactions between both predictors, we first conducted pairwise contrasts on both predictors comparing group differences between each cataract-reversal group and its matched control group across modalities and second pairwise contrasts testing for group differences separately for each modality condition.

Spearman’s rho, i.e., robust correlation coefficients were calculated between the two performance measures (bias and resolution) and CC-individuals’ key medical data (duration of visual deprivation, time period since visual restoration, and visual acuity). Partial correlations, correcting for the effect of age on TOJ performance ^9^, were used for temporal variables (duration of visual deprivation and time period since visual restoration). P-values were corrected for multiple comparisons using Benjamini and Hochberg’s procedure.

## Acknowledgments

Our work was supported by the European Research Council (ERC-2009-AdG 249425 CriticalBrainChanges) to BR and the German Research Foundation (DFG) with a research fellowship grant to SB (BA 5600/1-1) and Leibniz award money to BR (Ro 2625/10-1) as well as by the University of Hamburg with a post-doctoral fellowship to SB. We thank Marlene Hense, Nicola Kaczmareck, Carla Petroll, Lea Hornung, Deniz Froemke, and Rakesh Balachandar for help with data acquisition as well as Kabilan Pitchaimuthu for help with data curation.

## Author Contributions

Conceptualization, B.R.; Methodology, S.B., P.L., and B.R.; Software, S.B., and P.L.; Formal analysis, S.B.; Investigation, P.L., S.S.R., I.S., R.K., and B.R.; Data Curation, I.S.; Funding Acquisition, B.R.; Resources, B.R. and R.K.; Visualization, S.B.; Supervision, S.B. and B.R.; Writing – Original Draft, S.B. and B.R.; Writing – Approval and Edits, S.B., P.L., S.S.R., I.S., R.K., and B.R..

---

**Figure 1 – Figure Supplement 1.**
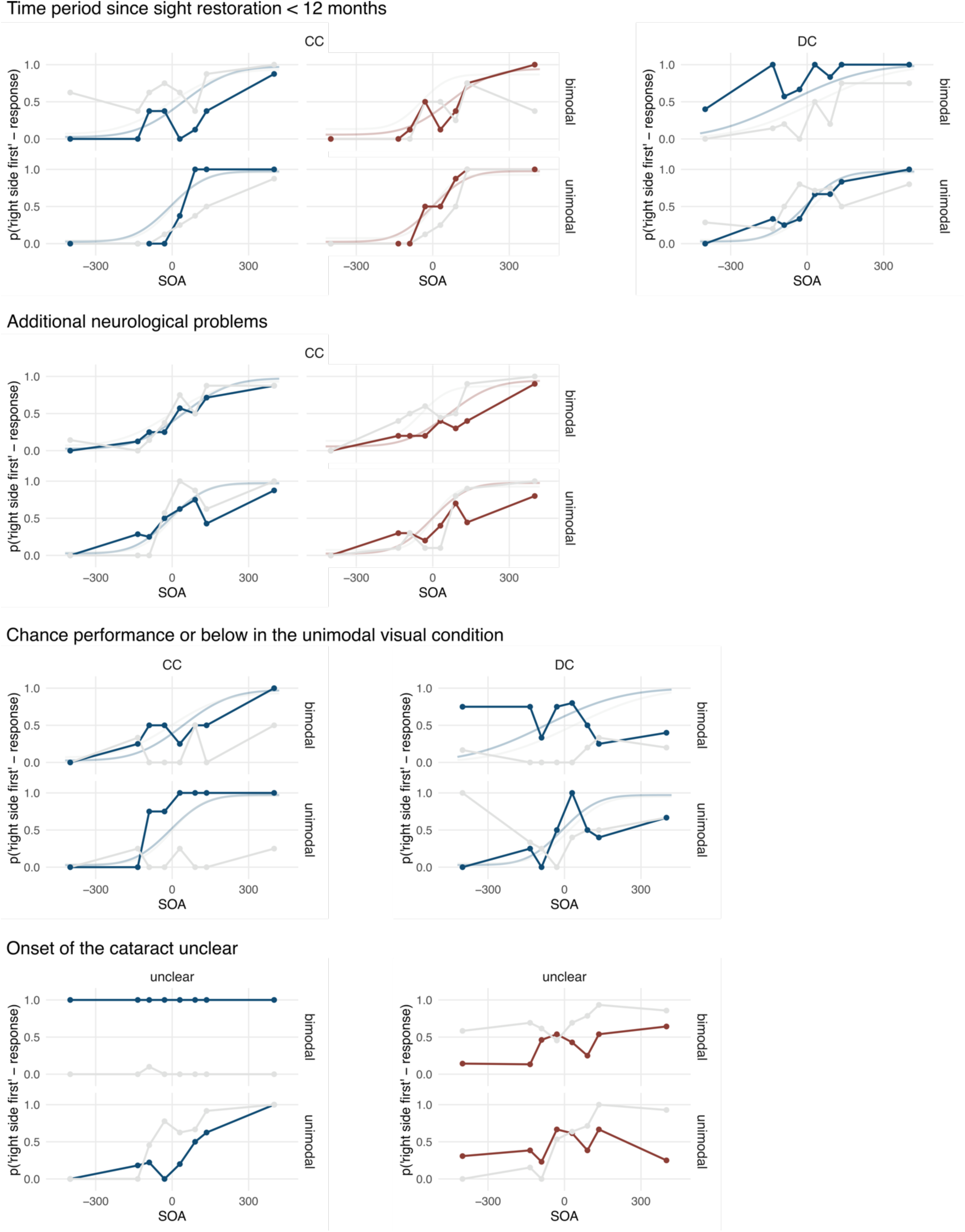
Excluded data. Group mean proportions of ‘right side first’-responses are shown as a function of the stimulus onset asynchrony (SOA) of the two stimuli, with negative values indicating ‘left side first’-stimulation. The data are split into responses to bimodal (top row) and unimodal (bottom row) stimulus pairs and according to the modality presented at the right side. Sigmoid curves fitted to the group mean data reported in Figure 1 are shown as a reference for participants for whom the onset of the cataract could be identified.

**Figure 1 – Figure Supplement 2.**
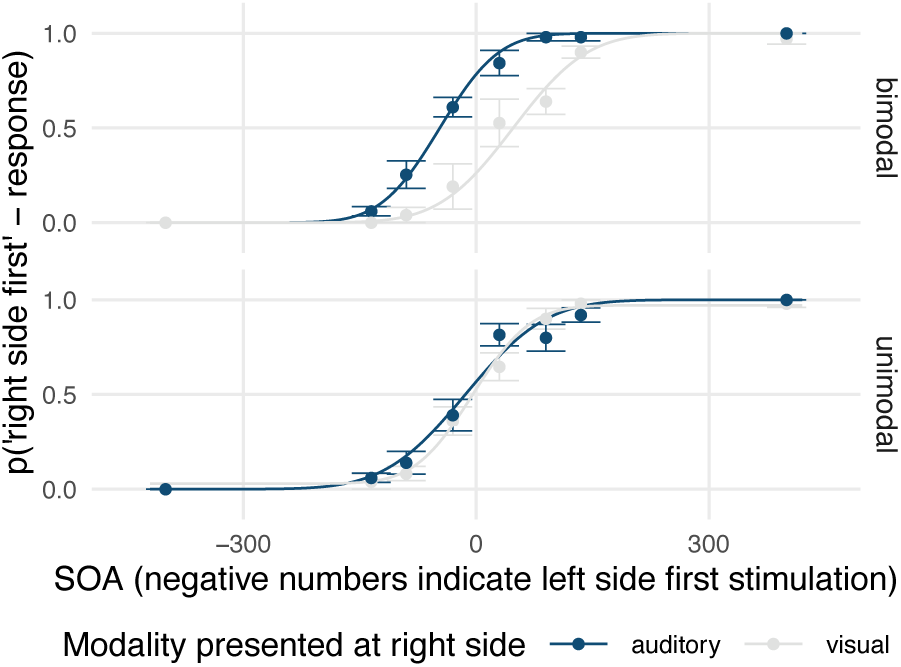
Visual-auditory temporal order judgments in typically sighted individuals in an additional experiment controlling for the effects of visual stimulus brightness. Five additional, typically sighted participants (all female and right-handed, 23-50 years old, mean age 34 years) completed the visual-auditory experiment. Here, a visual stimulus consisted of three activated LEDs whereas in the main experiment for typically sighted controls only one LED had been activated. Group mean proportions of ‘right side first’-responses are shown as a function of the stimulus onset asynchrony (SOA) of the two stimuli, with negative values indicating ‘left side first’-stimulation. The data are split into responses to bimodal (top row) and unimodal (bottom row) stimulus pairs and according to the modality presented at the right side. Participants showed a typical bias of perceiving visual stimuli as delayed compared to auditory stimuli.

